# Ellagic acid ameliorates atherosclerosis by targeting the epidermal growth factor receptor

**DOI:** 10.1101/2023.09.14.557846

**Authors:** Ye-Wei Huang, Li-Tian Wang, Huai-Liu Yin, Dan-Dan Hu, Jun Sheng, Xuan-Jun Wang

## Abstract

**BACKGROUND:** High levels of plasma low-density lipoprotein (LDL) are a key risk factor for atherosclerosis. Low-density lipoprotein receptor (LDLR) mediates the degradation of plasma LDL. Therefore, it may be possible to prevent and treat atherosclerosis by increasing the levels of LDLR. The natural polyphenolic compound ellagic acid (EA) has various biological activities. In mice, EA alleviated the progression of atherosclerosis; however, the underlying mechanism remains unclear.

**METHODS:** Molecular interaction, cell, and animal experiments were used to explore the role and mechanism of EA in improving atherosclerosis.

**RESULTS:** EA binds to the extracellular domain of the epidermal growth factor receptor (EGFR), thus activating the EGFR-extracellular signal-regulated kinase (EGFR-ERK) signaling pathway, stabilizing LDLR mRNA, and promoting the expression of LDLR protein. The development of EA-loaded human serum albumin nanoparticles enabled intravenous administration in animal experiments.

**CONCLUSIONS:** The research verified the in vivo effects of EA on the EGFR-ERK signaling pathway, LDLR levels, and atherosclerosis. EA may assist in the prevention and treatment of atherosclerosis.

---

Atherosclerosis is a chronic vascular disease caused by the deposition of cholesterol lipids in the inner wall of large and medium-sized arteries. It is characterized by high prevalence and associated with high disability and fatality rates. It is an important pathological basis for cardiovascular diseases, such as coronary heart disease, peripheral vascular disease, and arterial disease. Thus, it poses a serious threat to human life, health, and quality of life.^1,2^ Dyslipidemia has been recognized as the main cause of atherosclerosis by the medical community worldwide. Studies have shown that a decrease in low-density lipoprotein cholesterol (LDL-c) levels can effectively reduce the occurrence of atherosclerosis.^3^ Therefore, this approach has become the mainstay treatment for reducing the risk of atherosclerosis. At present, lipid-lowering drugs (e.g., statins) are the main agents used in the clinical treatment of atherosclerosis; however, the long-term use of these drugs is associated with serious adverse reactions, such as liver damage, skeletal muscle damage, and an increased risk of cancer. Consequently, the currently available countermeasures against atherosclerosis remain limited, and research and development on relevant drugs is of great significance.^4^

Low-density lipoprotein receptor (LDLR) is a key membrane receptor in the regulation of cholesterol homeostasis.^5^ LDL-c is mainly cleared from the blood circulation through LDLR-mediated endocytosis in the liver, which accounts for >70% of the total LDL particle clearance.^6–8^ Therefore, the role of LDLR in regulating LDL-c levels has attracted research attention in recent years. Messenger RNA (mRNA) turnover plays a key role in regulating protein levels, while adenosine and uridine-rich elements (AREs) of the untranslated region of mRNA (3’ untranslated region) control mRNA stability.^9^ LDLR mRNA is unstable, possibly due to rapid degradation after its interaction with ARE-binding proteins. Studies have shown that activation of epidermal growth factor receptor-extracellular signal-regulated kinase (EGFR-ERK) signaling can enhance LDLR mRNA stability. Moreover, ZFP36 ring finger protein like 1 (ZFP36L1) and ZFP36L2 control LDLR mRNA stability via the ERK-ribosomal protein S6 kinase (ERK-RSK) pathway.^10^ Berberine and 5-aztidine (5-Azac) can enhance the stability of LDLR mRNA by activating the ERK signaling.^11,12^ However, the precise mechanism underlying this post-transcriptional regulation remains unclear.

Ellagic acid (EA) is a natural polyphenolic compound that is widely present in various berries (e.g., pomegranate, blueberry, grape, strawberry) and tea.^13^ It has many biological activities (e.g., liver protection, lipid reduction, and cardiovascular disease improvement), and has demonstrated great potential in biomedicine.^14,15^ Studies have shown that EA can help to reduce the incidence of obesity. This is achieved by inhibiting the expression of key proteins of fat formation in fat precursor cells, as well as blocking the storage of triglycerides in fat cells to reduce lipid accumulation.^16^ EA is considered a potent naturally active compound owing to its non-toxic properties and multiple pharmacological actions. Nevertheless, it is almost insoluble in water (<10 μg/mL), has very low bioavailability (<0.06%), and has a blood concentration of only 0.1–0.4 μmol/L. Under the action of intestinal flora, the unabsorbed EA is mainly metabolized into urolithin A, B, C, D and a small part of urolithin M5, M6, etc.; hence its pharmacological action *in vivo* is greatly limited.^17^

Recently developed albumin-based drug delivery systems allow EA to be delivered by injection. As a natural protein, albumin is popular in the fields of drug delivery and disease treatment. Human serum albumin (HSA) is a single peptide composed of 585 amino acids, containing 17 disulfide bonds and one free sulfhydryl group. Its tertiary structure is crosslinked by three homologous domains (I, II, III) through disulfide bonds.^18^ Each domain is composed of two subdomains (A and B). There are two main drug-binding sites on subdomains IIA and IIIA, which can bind to a variety of hydrophobic drugs and improve their bioavailability.^19,20^ In addition, albumin is the most abundant protein in serum, with significant biocompatibility, biodegradability, non-immunogenicity, and other characteristics. Thus, it is an ideal carrier for the delivery of hydrophobic drugs, and is widely used in the pharmaceutical field.^21^

The US Food and Drug Administration has approved Abraxane (a nanoparticle of albumin-bound paclitaxel) for clinical use in the treatment of metastatic breast cancer.^22^ In addition, Yang et al. prepared HSA loaded with curcumin through the self-assembly method, which is also used for the treatment of various diseases.^23^ There are numerous functional carboxyl groups and amino groups on the surface of HSA, which have the ability to form covalent bonds with compounds.^24^ In addition, albumin is widely used for the production of hydrophobic drug carriers owing to its simple preparation and high yield.

In this study, HepG2 cells were used to conduct *in vitro* studies. The objective was to investigate the effect of EA on LDLR expression and the molecular mechanism involved in this process. EA-loaded HSA nanoparticles (EA-naps) were prepared through the self-assembly method using a wide range of sources of HSA as drug carriers to improve the safety profile of the preparation for intravenous injection. Furthermore, in vivo experiments were conducted in apolipoprotein E (ApoE)^-/-^ mice to explore the efficacy and mechanism of EA in the treatment of atherosclerosis.

## MATERIALS AND METHODS

### Materials

EA (purity: >98%) was purchased from Shanghai Yuanye Biotech. Co. Ltd (Shanghai, China). HSA (purity: >98%; Vetec™ reagent grade), MTT, DTT, Dil-LDL and improved oil Red O staining kit were obtained from Solarbio (Beijing, China). SYBR Green real-time PCR Master Mix was purchased from Takara (Beijing, China). Other antibodies included anti-LDLR (ET1606, Huabio, Hangzhou, China), anti-EGFR (2085S, Cell Signaling Technology, MA, USA), anti-phospho-EGFR (48576SF, Cell Signaling Technology, MA, USA), anti-ERK1/2 (4696S, Cell Signaling Technology, MA, USA), anti-phospho-ERK1/2 (4370S, Cell Signaling Technology, MA, USA), and anti-β-actin (66009-1-IG, Proteintech, Wuhan, Hubei, China). Male C57BL/6 ApoE^-/-^ mice (age: 6 weeks, weight: 18–20 g) were purchased from Changzhou Card Vince Laboratory Animal Co. Ltd., (Changzhou, China; license number: SCXK(SU)2016-0010). HepG2 cells were obtained from American Type Culture Collection (Manassas, VA, USA). The recombinant EGFR extracellular domain (catalog number: 10001-H08H) was obtained from Sino Biological (Beijing, China).

### Preparation of EA-naps

EA-naps were prepared through the self-assembly method.^44^ Specifically, HSA (40 mg) was dissolved in tris buffer (40 mL, pH 7.4, 5 mM) at 40℃, followed by the addition of DTT (final concentration: 5 mM) was added; the solution was stirred at 40℃ for 5 min. Next, under continuous stirring, EA (4 mg) (dimethyl sulfoxide was used as solvent) was added, and the solution was thoroughly mixed for 10 min. Ultrasound was utilized at low power (300 W) for 3 min to disperse the aggregated particles. EA-naps were extensively dialyzed using a dialysis bag (molecular weight: 8 kDa) to remove any residual DTT and free EA.

### Morphology, particle size, dispersion index, and Zeta potential

The morphologies of EA-naps were determined using a JEM-2100 high-resolution transmission electron microscope (JEOL). The solution was placed on a 200-mesh grid composed of carbon-coated copper and negatively stained with phosphotungstic acid. Subsequently, the particles were observed under a scanning electron microscope. The powder sample was placed on a silicon wafer, sputtered with platinum, and examined through scanning electron microscopy. The particle size, polymer dispersion index, and Zeta potential were determined using a Malvern Laser scattering particle size analyzer (Malvern Instruments, Malvern, UK) at room temperature.

### Encapsulation rate and loading rate of EA-naps

In C18 column high-performance liquid chromatography (HPLC; LC-2010 CHTSystem; Shimadzu, Kyoto, Japan): the dimensions of the column were 250 mm × 4.6 mm, 5 μm; the mobile phase consisted of acetonitrile:0.03% trifluoroacetic acid (15:85; volume/volume [v/v]); the flow rate was 1.0 mL/min; the column temperature was 30℃; the detection wavelength was 254 nm; and the sample size was 10 μL. This method was used to determine the concentration of the drug. EA-nap suspension (1 mL) was placed into an ultrafiltration centrifuge tube (interception molecular weight: 8 kDa; Millipore, Ireland), and centrifuged at −4℃ for 20 min at 12,000 r/min. The filtrate was obtained, and the levels of free EA (M_free_) were determined through HPLC. Suspension (1 mL) was taken, methanol precipitated protein (3 mL) was added, and the nanoparticles were destroyed through ultrasonication for 10 min. The filtrate was filtered through microporous membrane (pore size: 0.22 μm), and the total EA (M_total_) was determined using HPLC.

The encapsulation rate (EE%) and drug loading rate (LC%) were calculated as follows:

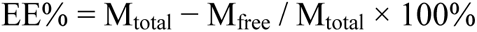

M_total_ is the amount of total EA, and M_free_ is the amount of EA that remained in the supernatant.

The encapsulation rate was calculated using the following formula:

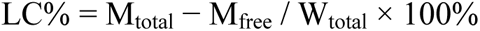

M_total_ is the amount of total EA, and W_total_ is the weight of lyophilized nanoparticles.

### Coomassie Brilliant Blue staining

Sodium dodecyl sulfate-polyacrylamide gel electrophoresis was performed as previously described.^44^ In a vertical gel electrophoresis device (Bio-Rad, USA), 4% concentrated gel and 8% separated gel were used. The samples (2 μg/μL) were mixed with loading solution with or without β-mercaptoethanol, and boiled at 95℃ for 10 min. The prepared samples were added to the gel, and electrophoresis was performed for 80 min at 120V at constant pressure. Next, the gel was stained with Coomassie Brilliant Blue R-250 for 20 min and washed in decolorizing solution. Finally, the protein bands on the gel were visualized using the ChemiDoc imaging system (Bio-Rad).

### In vitro release experiment

The effect of drug release by EA-naps was measured using the dialysis bag method. After shaking and mixing, the prepared EA-naps were placed into a dialysis bag (interception molecular weight: 8 kDa), and dissolution medium (3 mL) was added. The dissolution medium was phosphate-buffered saline (PBS) containing 0.1 v/v Tween 80 (pH 7.4) (200 mL), the rotational speed was 100 rpm/min, and the temperature was maintained at 37±0.5℃. Samples (0.5 mL) were obtained at specific time points (i.e., 0, 2, 4, 8, 12, and 24 h); the volume was adjusted with fresh buffer solution to account for sampling. Finally, the levels of drug were determined by HPLC.

### MTT experiment

HepG2 cells were cultured in high-glucose medium supplemented with 10% fetal bovine serum and 1% antibiotics. The cells were maintained in a humid atmosphere at 37℃ with 5% CO_2_. MTT assay was used to evaluate cell viability and, thus, determine the safety profile of EA. HepG2 cells were treated with different concentrations of EA for 24 h, MTT solution (20 μL, 5 mg/mL) was added, and the cells were incubated at 37℃ for 4 h. Thereafter, the culture medium was discarded and dimethyl sulfoxide (100 μL) was added to dissolve purple crystals in each well. The absorption was measured at 492 nm wavelength using a microplate reader (Synergy4, multi-mode microplate reader; BioTek, Winooski, VT, USA).

### Western blotting

For the extraction of total protein, cell lysis solution was added to HepG2 cells and liver tissues obtained from ApoE^-/-^ mice. The protein content was detected using bicinchoninic acid working solution (Biyuntian), and the sample volume was calculated. The protein samples were mixed with the carrier solution and boiled at 95℃ for 10 min to denature the protein. Subsequently, the samples were placed into the injection hole of the rubber plate. After sodium dodecyl sulfate-polyacrylamide gel electrophoresis, the glue protein was transferred to a polyvinylidene difluoride membrane. Next, the membrane was blocked with 5% skim milk powder for 1 h, and incubated with rabbit anti-LDLR antibody on a shaking platform at 4℃ overnight. The next day, the membrane was incubated with goat anti-rabbit IgG secondary antibody on a shaking platform at room temperature for 1 h. Target protein blotting images were obtained using enhanced chemiluminescence mixed luminescent solution on the imaging system (Media Cybernetics, Inc.).

### Real-time PCR

Total RNA was extracted from HepG2 cells and liver tissues using Trizol reagent. The total RNA was reversely transcribed into cDNA with SYBR Green real-time PCR master mix, and PCR was performed according to the instructions provided by the manufacturer. The conditions for the real-time quantitative PCR assay were as follows: predenaturation at 95℃ for 30 s; denaturation at 95℃ for 5 s; and annealing at 60℃ for 30 s; for a total of 40 cycles. Subsequently, the levels of PCR products were normalized to those of β-actin. The cycle threshold (Ct) contrast method (2^-ΔΔCt^) was used to determine the relative expression of each target gene.

### Dil-LDL uptake experiment

HepG2 cells were treated with EA for 20 h; subsequently, Dil-LDL (20 μg/mL) was added, and the cells were stored at 37℃ in the dark. Thereafter, the cells were washed thrice with PBS and fixed with 4% paraformaldehyde for 20 min at room temperature. Finally, the cells were sealed with an anti-fluorescence quencher containing 4’,6-diamidino-2-phenylindole, and observed and photographed under a fluorescence microscope.

### SPR analysis

The Biacore S200 instrument (GE Healthcare, Uppsala, Sweden) was used to determine the interaction between proteins and small molecules based on the principle of SPR. The purified EGFR recombinant extracellular domain was first coupled to the surface of a CM5 sensor chip (GE Healthcare Biosciences)). Kinetics and affinity analyses of EA injected into the EGFR fixed chip sensor surfaces were performed using PBS + 0.05% Tween 20 (v/v) buffer.

### Molecular docking experiment

Molecular docking of EA to EGFR was performed using AutoDock (v.4.2) version. The EGFR (PDB ID: 3njp) protein structure was downloaded from the PDB database^27^. EGFR-related parameters were set as follows: center_x = 11.5, center_y = 20.5, center_z = 18.1; Search Spaces: size_x: 50, size_y: 50, size_z: 50 (spacing of 0.375 A between each square point), and tiveness: 10. The remaining parameters were maintained at the default values. Tiveness: 50 three-dimensional structures were downloaded in Scientific Data Format from the PubChem database according to Chemical Abstracts Service number of EA, import the structure was imported into ChemBio3D Ultra 14.0 for energy minimization, and the Minimum RMS Gradient was set at 0.001, small molecules were saved in mol2 format, and optimized small molecules were saved in “pdbqt” format. Conduct motion pattern analysis was conducted, and graphs were produced.

### Grouping and treatment of animals

C57BL/6J ApoE^-/-^ mice (male; age: 6 weeks) were purchased from Changzhou Card Vince Laboratory Animal Co., Ltd. (license number: SCXK(SU)2016-0010). The animals were housed in an environment with a constant temperature of 18–24℃ and humidity of 40–70% on a 12-h day/night cycle, with free access to food and water. All experiments involving animals were performed in accordance with the Guidelines for The Care and Use of Experimental Animals of Yunnan Agricultural University and approved by the Animal Ethics Committee of Yunnan Agricultural University (approval number: YNAU202103042). There were no adverse events observed. ApoE^-/-^ mice were randomly divided into three groups (n = 7 per group), namely the LFD group, HFD group, and EA-naps group. The LFD group was fed an ordinary diet (article number: HFHC; Deitz), while the other groups were fed a HFD (article number: ASLF4; Deitz). In the EA-naps group, EA (concentration: 25 mg/mL) was intravenously injected into the mice. In the control and HFD groups, the mice received intravenous injections with an equal volume of normal saline once daily for 12 consecutive weeks. Thereafter, the mice were weighed and monitored on a weekly basis. After 12 weeks, the ApoE^-/-^ mice were sacrificed, and liver and aorta tissues were collected. Blood was obtained from mouse eyeballs, the samples were centrifuged at 3,500 r/min for 15 min, serum was separated, and the samples were stored at −80℃. Liver tissues were placed in liquid nitrogen and tissue fixation solution, and aorta tissues were placed in tissue fixation solution.

### Hematoxylin-eosin staining and immunohistochemistry

Liver and aorta samples were immobilized in tissue fixator and embedded in paraffin. The tissue samples were sliced (thickness: 5 μm), stained with hematoxylin and eosin (Solarbio, Beijing, China), sealed with gum, and finally observed and photographed under a Leica DM2500 microscope (Wetzlar, Germany). Next, immunohistochemical staining was performed. The slices were soaked in a sodium citrate antigen-repair buffer, placed in the microwave for repair, and cooled to room temperature. Next, the slices were incubated with 10% goat serum at room temperature for 30 min to prevent non-specific binding of antibodies. The sections were incubated with primary antibodies against LDLR (1:200), phosphorylated-EGFR (p-EGFR; 1:200), αSMA secondary antibody (1:200), and CD68 (1:200) at 4℃ overnight. The next day, the slices were incubated with secondary antibody at room temperature for 30 min, stained brown with DAB color developing solution, and reversed stained with hematoxylin. Finally, the slices were sealed with neutral gum. The sections were observed and photographed under an optical microscope (Eclipse Ti-E; Nikon, Tokyo, Japan).

### Oil Red O staining

Liver, gross aorta, and aortic sinus tissues obtained from ApoE^-/-^ mice were stained using a modified oil Red O kit. These tissues were immersed in 60% isopropyl alcohol solution for 5 min, and then in oil Red O staining solution for 10 min. The differentiation was differentiated by immersion in 60% isopropyl alcohol solution for 2–3 s and terminated by rinsing with tap water. The liver and aortic sinus tissues were sealed with glycerin gelatin. Finally, the liver was observed and photographed under a Leica DM2500 microscope (Wetzlar, Germany).

### EVG staining

According to EVG- Verhöeff staining kit (G1597, Solarbio, China). Aortic sinus tissue sections obtained from ApoE^-/-^ mice were soaked in EVG dye solution for 30 min and rinsed with tap water. The ferric chloride solution was differentiated, and the background color and elastic fiber color were observed under a under a Leica DM2500 microscope (Wetzlar, Germany). The elastic fiber color was purple-black.

### Statistical analysis

All measurement data were expressed as the mean + standard. All statistical analyses were performed using GraphPad Prism 8.0 (GraphPad Software Inc., San Diego, CA, USA). Student’s *t*-test was used for the statistical analyses, and P-values <0.05 denoted statistically significant differences.

## RESULTS

### EA promoted LDLR expression in HepG2 cells

In this study, 3-(4,5 dimethyl-thiazol-2,5 biphenyl) tetrazolium bromide (MTT) assay was used to evaluate whether EA concentrations <40 μM exerted a significant effect on the viability of HepG2 cells (Figure 1A). Since LDLR plays a key role in the occurrence and progression of atherosclerosis, we evaluated the effect of EA on LDLR protein levels. The results showed that EA significantly increased LDLR protein levels in HepG2 cells in a dose-dependent manner (Figure 1B, C). We also investigated the effect of EA on the uptake of Dil-LDL by HepG2 cells, observing that EA promoted LDLR function on the cell membrane surface (Figure 1D). These results suggest that EA can induce an increase in LDLR protein levels, thereby promoting LDL uptake by HepG2 cells.

**Figure 1.**
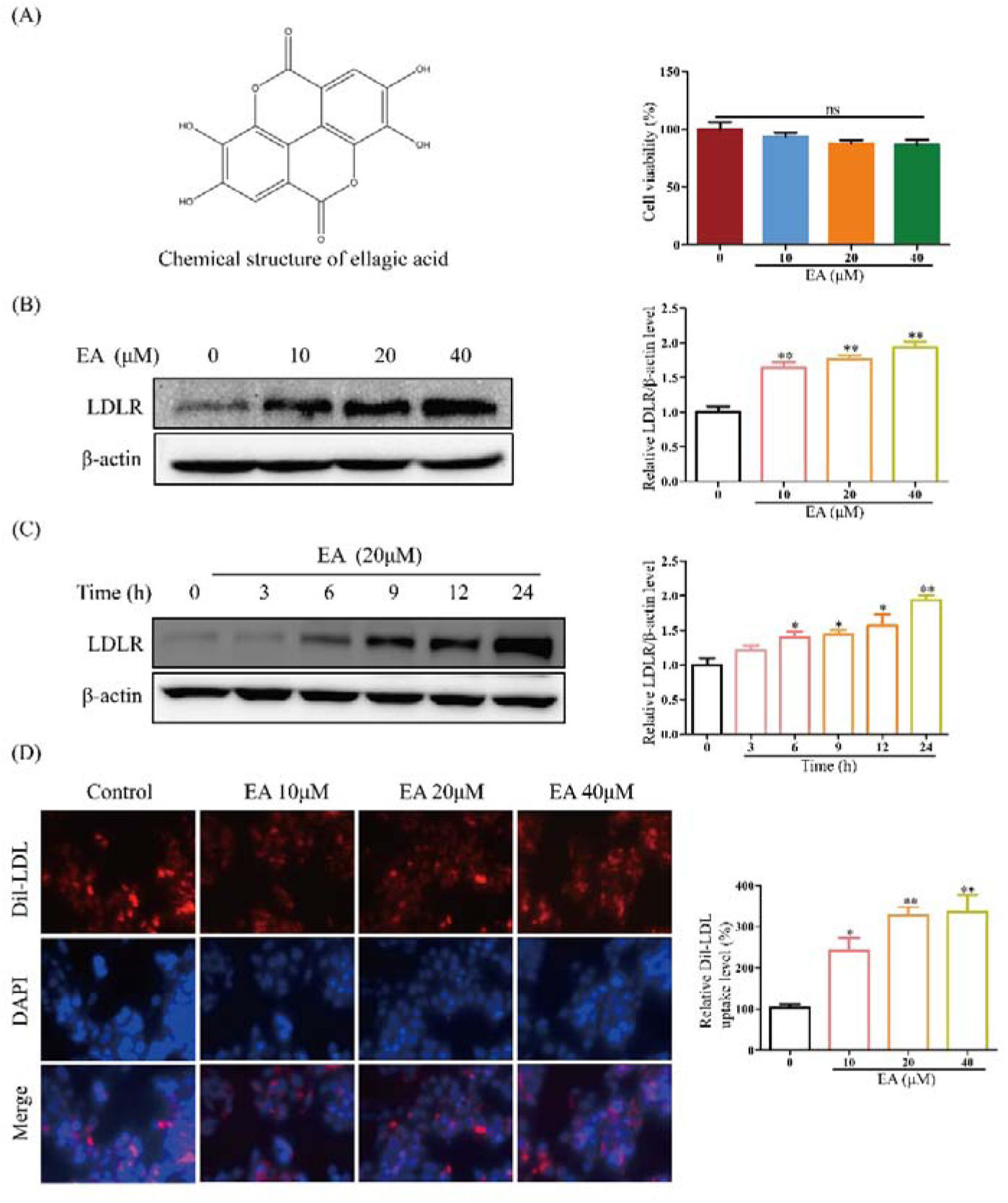
EA induced an elevation in LDLR protein levels and promoted LDL uptake by HepG2 cells. (A) Cytotoxicity of EA (10, 20, and 40 μM) in HepG2 cells. (B) Effects of different concentrations of EA (10, 20, and 40 μM) on LDLR expression in HepG2 cells. (C) Effect of EA (20 μM) on LDLR expression in HepG2 cells at different treatment times (3, 6, 9, 12, and 24 h). (D) Fluorescence microscopy was used to observe the uptake of Dil-LDL by EA-treated HepG2 cells for 20 h, and quantitative analysis of Dil-LDL uptake. Data represent the mean ± SEM (n = 3). *P < 0.05 and **P < 0.01. EA, ellagic acid; LDL, low-density lipoprotein; LDLR, low-density lipoprotein receptor; SEM, standard error of the mean.

### EA enhanced LDLR mRNA stability by activating the EGFR-ERK signaling pathway

We examined the effect of EA on LDLR mRNA levels to explore the mechanism through which it induces an increase in LDLR protein expression. The results showed that EA induced a significant increase in LDLR mRNA levels in HepG2 cells (Figure 2A). Increased expression of LDLR mRNA can be achieved by enhancing its stability. Hence, we further evaluated the effect of EA on LDLR mRNA stability using actinomycin D. The results showed that EA significantly improved the stability of LDLR mRNA by extending its half-life from approximately 1.5 h to 2.3 h (Figure 2B).

**Figure 2.**
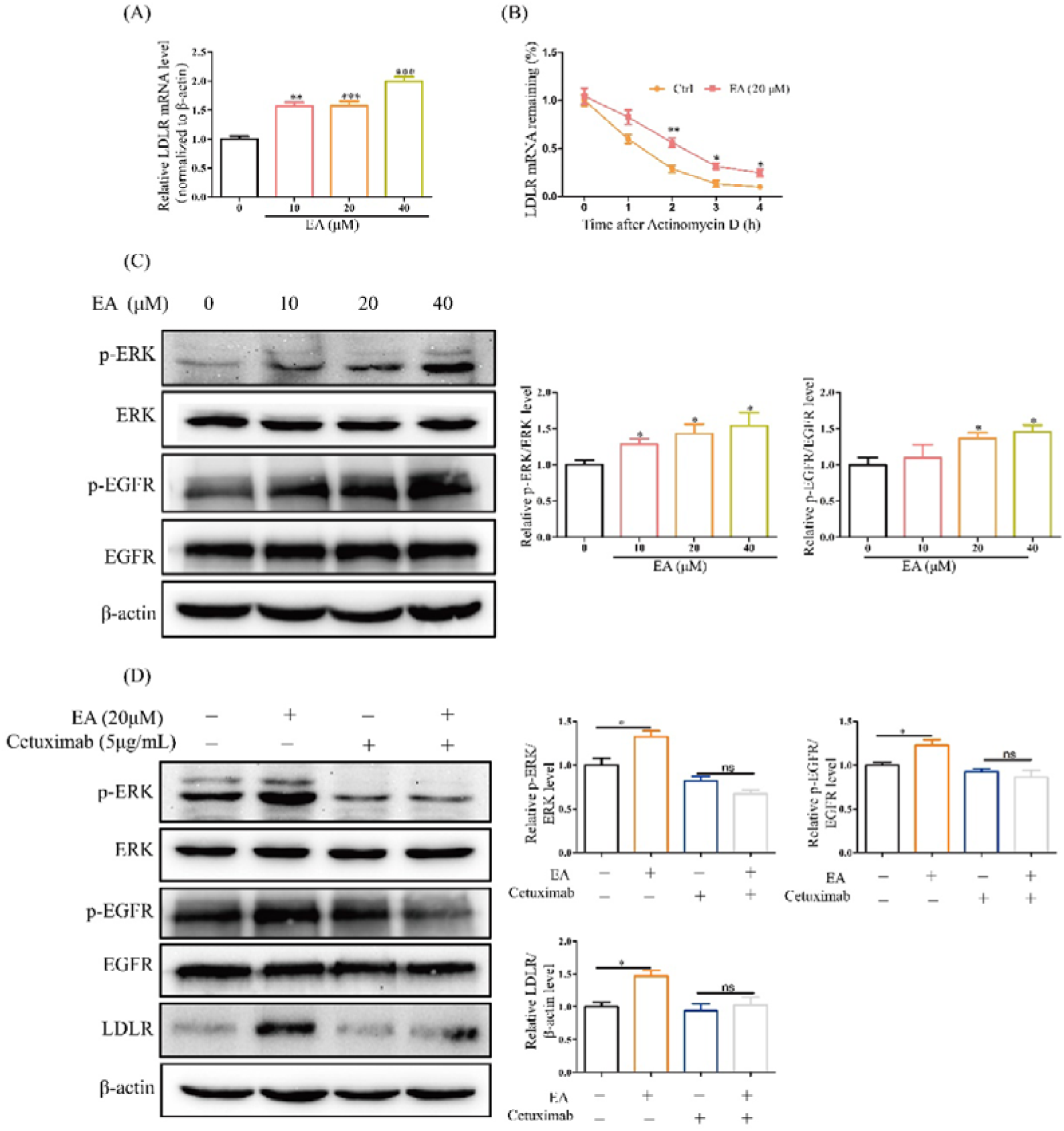
EA increased LDLR expression through the EGFR-ERK signaling pathway. (A) Effect of EA (treatment for 24 h) on LDLR mRNA levels in HepG2 cells. (B) The effect of EA on the stability of LDLR mRNA, observed under the mRNA synthesis of HepG2 cells was inhibited by actinomycin D. (C) Effects of different concentrations of EA (10, 20, and 40 μM) on p-ERK and p-EGFR levels in HepG2 cells. (D) Effects of EA on p-ERK, p-EGFR, and LDLR levels after treatment with cetuximab to block the EGFR pathway in HepG2 cells. Data represent the mean ± SEM (n = 3). *P < 0.05, **P < 0.01, and ***P < 0.001. EA, ellagic acid; EGFR, epidermal growth factor receptor; ERK, extracellular signal-regulated kinase; LDLR, low-density lipoprotein receptor; p-EGFR, phosphorylated-EGFR; p-ERK, phosphorylated-ERK; SEM, standard error of the mean.

Studies have shown that EGFR is closely related to the metabolic pathway of cholesterol, and activation of EGFR-ERK signaling pathway can lead to increased mRNA levels of LDLR.^12,25,26^ We also sought to investigate whether the EGFR-ERK signaling pathway is involved in the stabilizing effect of EA on LDLR mRNA. Thus, we determined the effect of EA on the EGFR-ERK signaling pathway. The results showed that treatment with EA significantly increased the phosphorylation levels of EGFR and ERK in HepG2 cells (Figure 2C). In addition, our experimental results showed that the activation of EGFR-ERK signaling by EA could be blocked by cetuximab, a monoclonal antibody against EGFR. Cetuximab also significantly suppressed the increase in LDLR protein levels induced by EA (Figure 2D). These results indicate that EA exerts a role in increasing the stability of LDLR mRNA by acting on EGFR.

### EA directly bound to the extracellular domain of EGFR

To further verify whether EA can promote LDLR expression by directly acting on EGFR, we assessed the interaction between EA and EGFR extracellular segments using surface plasmon resonance (SPR) assay and computer simulated molecular docking techniques. The results showed that EA can directly bind to the extracellular domain of EGFR (dissociation constant [KD] = 4.33E-7M) (Figure. 3A). AutoDock (v 4.2) was used to conduct molecular docking tests between EA and EGFR (Protein Data Bank identifier [PDB ID]: 3njp).^27^ The results showed that the binding energy was −7.1 kcal/mol (Figures. 3B–D). The hydroxyl group of EA can form hydrogen bonds with the B chains GLU221, B chains HIS209, A chains GLU221, A chains THR239, and ASP238 of EGFR protein. The lengths of these hydrogen bonds are 2.07A, 1.93A, 2.21A, 2.6A, and 2.27A, respectively. This shows that hydrogen bonds play an important role in the binding between EA and EGFR (Figures. 3E, F). In addition, residues of HIS209, GLU211, CYS236, LYS237 of chain A, and LYS237 of chain B of EGFR were involved in the formation of hydrophobic interactions. These hydrogen bonds and hydrophobic interactions result in the relatively stable binding of EA to the “pocket” position of EGFR protein, which plays an important role in the structural stability of the EGFR-EA complex. These results suggest that EA can activate the EGFR-ERK signaling pathway by acting directly on the extracellular segment of EGFR, thus promoting LDLR expression.

**Figure 3.**
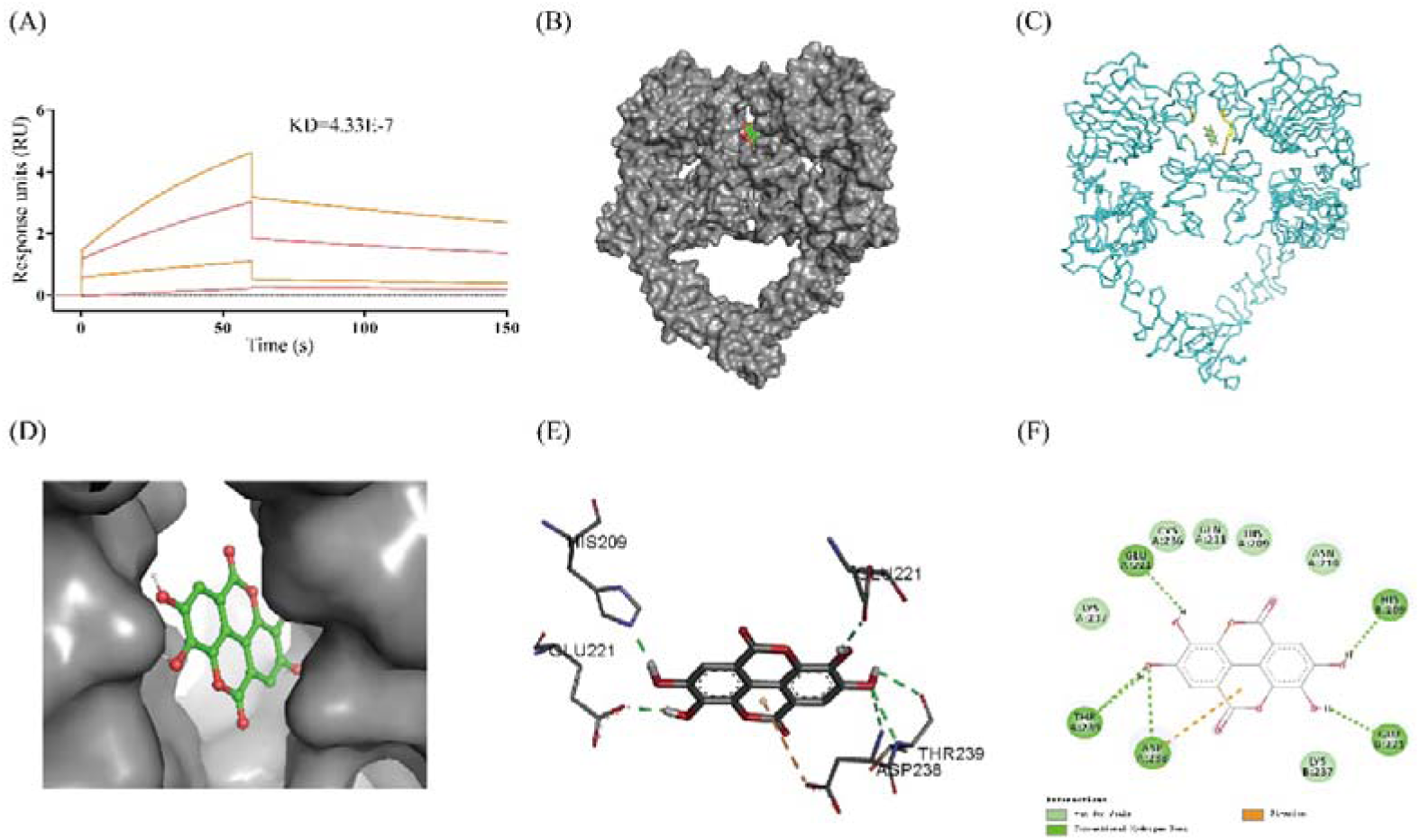
EA directly bound to the extracellular domain of EGFR. (A) Binding of EA and EGFR (extracellular domain). (B) Computer simulation 3D image of EA docking with the extracellular domain of EGFR. (C) 3D docking diagram of EA binding to the extracellular segment of EGFR. (D) EA zoom-in binding pocket. (E) 3D model of the interaction between EA and EGFR. Green and yellow dotted lines indicate hydrogen bond and hydrophobic interactions, respectively. (F). 2D model of the interaction between EA and EGFR. 2D, two-dimensional; 3D, three-dimensional; EA, ellagic acid; EGFR, epidermal growth factor receptor; SPR, surface plasmon resonance.

### EA-naps activated the EGFR signaling pathway and increased LDLR levels in HepG2 cells

The chemical properties of EA are responsible for its poor solubility in water or alcohol. We prepared EA-naps through the self-assembly method to improve the solubility and bioavailability of EA (Figure 4A). The albumin used in the EA-naps is widely used in the delivery of various hydrophobic therapeutic drugs owing to its favorable safety and biocompatibility. This lays a solid foundation for the successful verification of the role and molecular mechanism of EA in improving atherosclerosis in animal experiments. Subsequently, we tested whether EA-naps had equivalent biological activity to that of EA in HepG2 cells. The results showed that, similar to EA, EA-naps can activate the EGFR-ERK signaling pathway and increase the levels of LDLR (Figure 4B, C).

**Figure 4.**
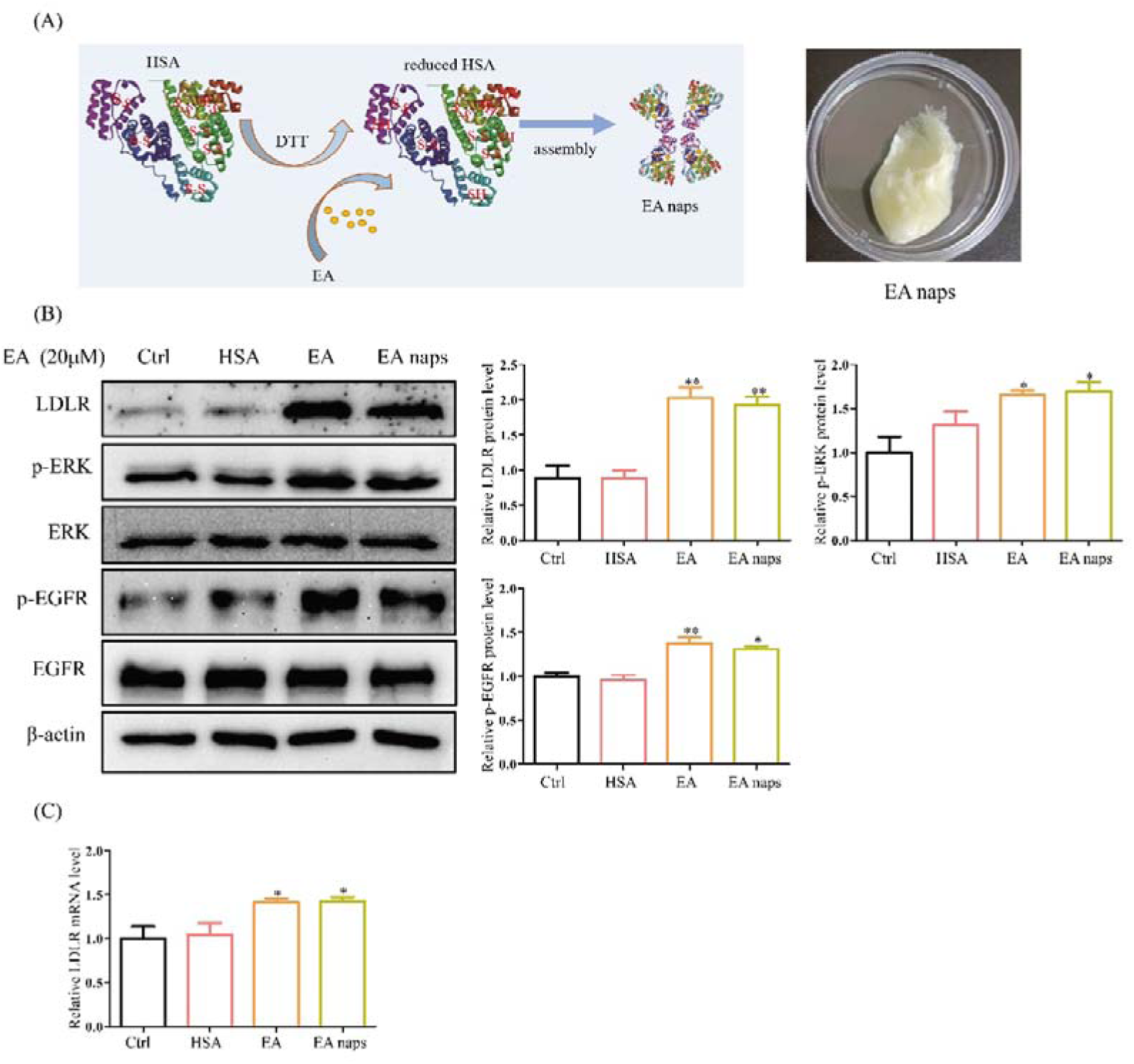
EA-naps increased the levels of LDLR in HepG2 cells. (A) Diagram of the EA-naps composite and appearance of the EA-naps powder. (B) Protein levels of LDLR, p-ERK, and p-EGFR in HepG2 cells treated with EA or EA-naps. (C) mRNA levels of LDLR in HepG2 cells treated with EA or EA-naps. Data represent the mean ± SEM (n = 3). *P < 0.05 and **P < 0.01. EA, ellagic acid; EA-naps, ellagic acid-loaded human serum albumin nanoparticles; EGFR, epidermal growth factor receptor; ERK, extracellular signal-regulated kinase; HSA, human serum albumin; LDLR, low-density lipoprotein receptor; p-EGFR, phosphorylated-EGFR; p-ERK, phosphorylated-ERK; RT-PCR, real-time polymerase chain reaction; SEM, standard error of the mean.

### EA-naps significantly improved atherosclerosis in ApoE^-/-^ mice

We sought to verify *in vivo* that EA enhances LDLR expression by activating the EGFR-ERK signaling pathway, thus improving atherosclerosis. For this purpose, we used a C57BL/6J ApoE^-/-^ mouse model of atherosclerosis to explore the anti-atherosclerotic effect and mechanism of EA-naps. The concentration of EA in mouse plasma reached 6.29 μmol/L.

The results of oil Red O staining analysis on gross aorta and aortic sinus tissues obtained from mice showed that the atherosclerotic plaque area in the high-fat diet (HFD) group was significantly increased compared with that observed in the low-fat diet-fed (LFD-fed) group. Notably, treatment with EA-naps significantly reduced the plaque deposits in the gross aorta and aortic sinus of HFD-fed mice (Figure 5A, B).

**Figure 5.**
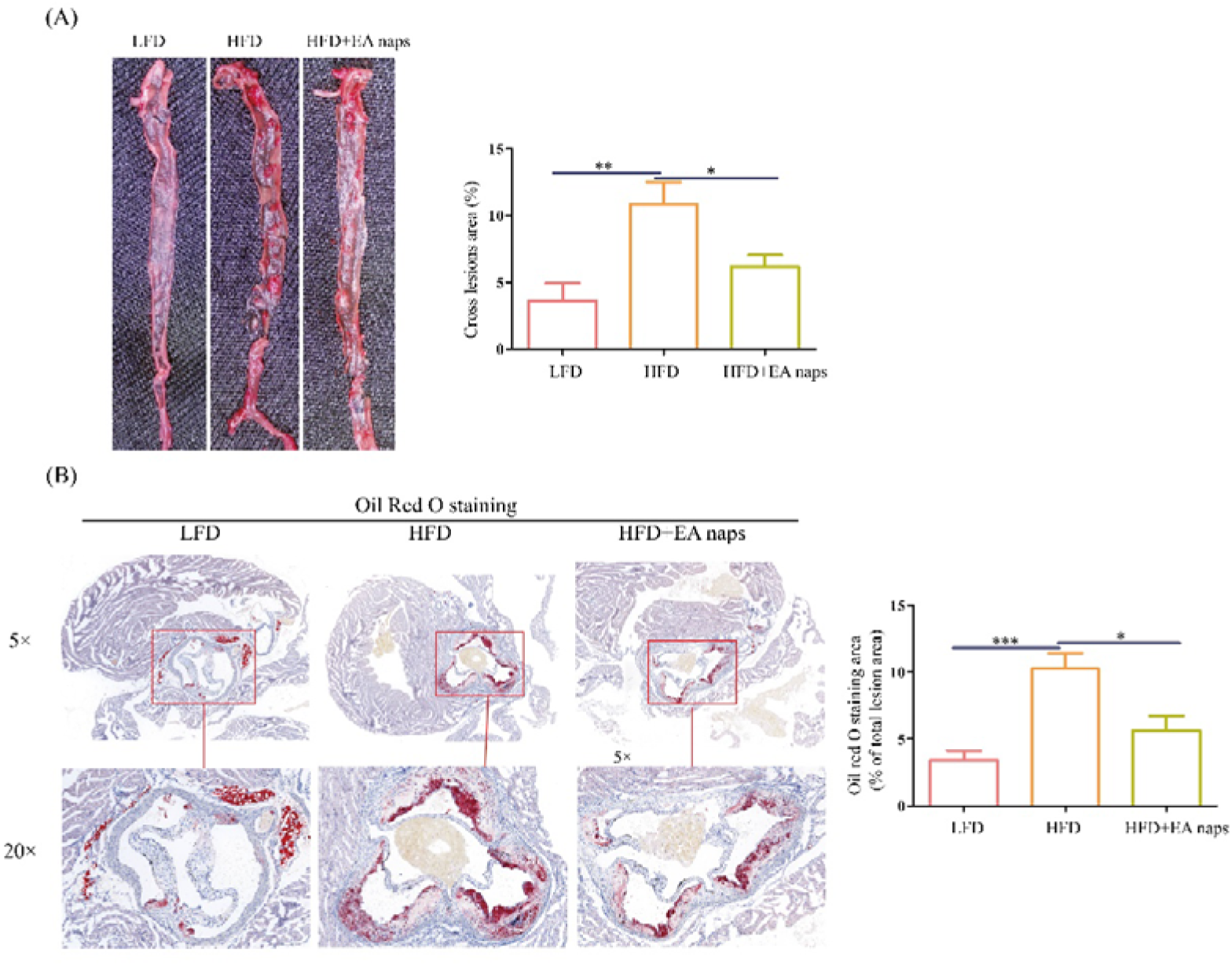
EA-naps ameliorated atherogenesis in HFD-fed ApoE^-/-^ mice. (A) Atherosclerotic plaques in the whole aorta of mice were stained with oil Red O, and statistical analysis of the stained plaque areas in the whole aorta. (B) The aortic root of mice was determined by oil Red O staining, and the stained plaque areas were quantified using ImageJ software. *P < 0.05, **P < 0.01, and ***P < 0.001. Data represent the mean ± SEM of six biological replicates. LFD group: chow diet; HFD group: high-fat and high-cholesterol diet, and treatment with uncoated EA nanoparticles; HFD+EA-naps group: high-fat and high-cholesterol diet, and treatment with coated EA nanoparticles. ApoE, apolipoprotein E; EA, ellagic acid; EA-naps, ellagic acid-loaded human serum albumin nanoparticles; HFD, high-fat diet; LFD, low-fat diet; SEM, standard error of the mean.

The occurrence and development of atherosclerosis include infiltration by macrophages from the lumen, migration of vascular smooth muscle cells (VSMCs) from the media into the intima, and reduction of the elastic fibers in the aortic root.^28^ The results of immunofluorescence staining for macrophage marker CD68 showed that EA-naps significantly reduced the levels of CD68 in the aortic roots of HFD-fed ApoE^-/-^ mice. Moreover, macrophage infiltration in atherosclerotic plaques was effectively alleviated by treatment with EA-naps (Figure 6A). The results of immunofluorescence staining for alpha-smooth muscle actin (αSMA) (a marker of VSMCs) showed that EA-naps could significantly reduce the levels of VSMCs in the aortic root of HFD-fed ApoE^-/-^ mice (Figure 6B). Elastic Verhoeff-Van Gieson (EVG) staining analysis showed that EA-naps significantly increased the amount of elastic fibers in the aortic roots of HFD-fed ApoE^-/-^ mice (Figure 6C). Hematoxylin-eosin staining of the aortic root revealed that EA-naps could significantly reduce the necrotic area of the aortic root in HFD-fed ApoE^-/-^ mice (Figure 6B). These results indicate that EA-naps are effective in alleviating the occurrence and development of atherosclerosis.

**Figure 6.**
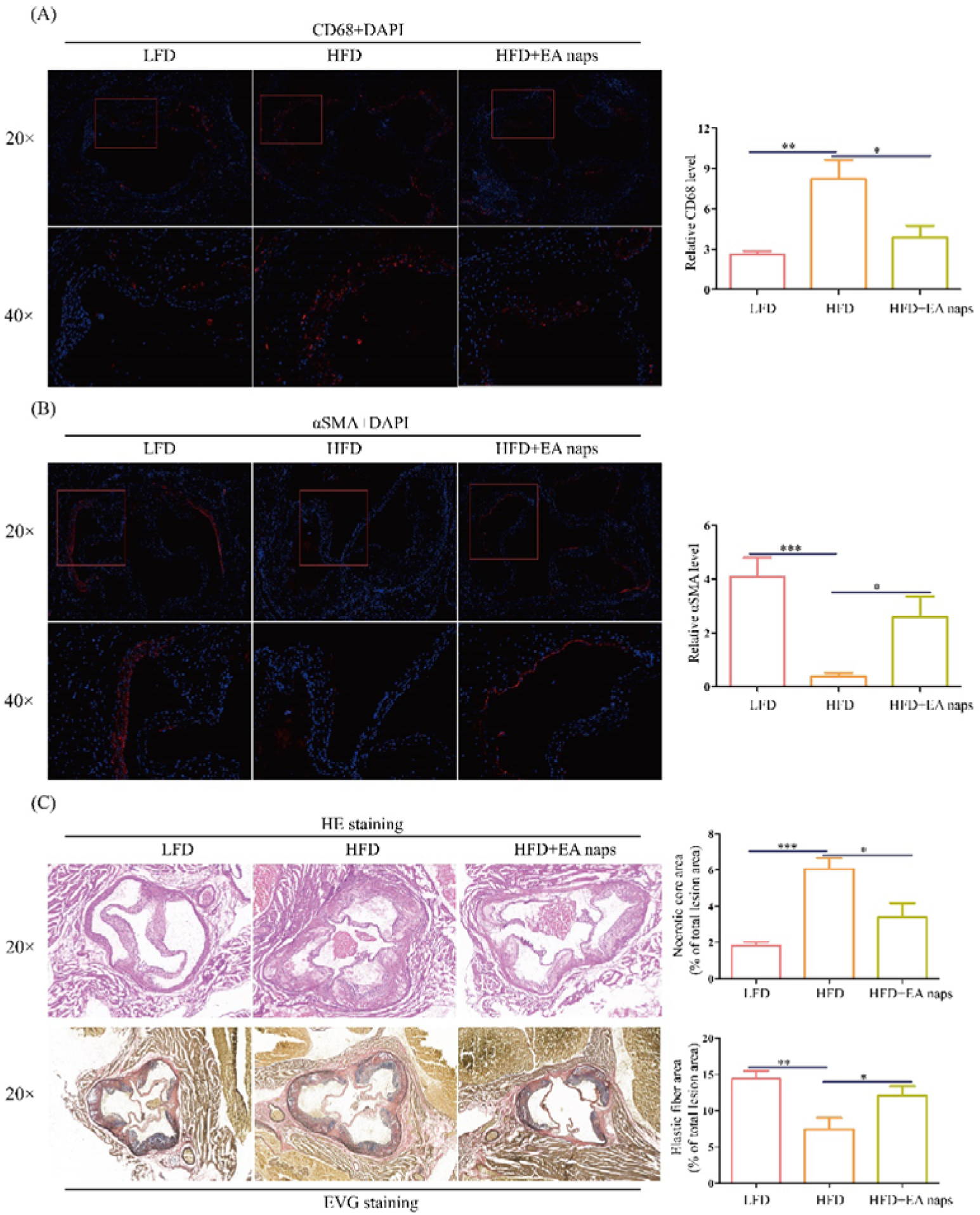
EA-naps ameliorated atherogenesis in HFD-fed ApoE^-/-^ mice. Immunohistochemical images of aortic root atherosclerotic lesions of mice stained for CD68 (A) or αSMA (B). (C) Aortic root atherosclerotic lesions of mice stained for HE or EVG. *P < 0.05, **P < 0.01, and ***P < 0.001. Data represent the mean ± SEM of six biological replicates. LFD group: chow diet; HFD group: high-fat and high-cholesterol diet, and treatment with uncoated ellagic acid nanoparticles; HFD+EA-naps group: high-fat and high-cholesterol diet, and treatment with coated ellagic acid nanoparticles. ApoE, apolipoprotein E; αSMA, alpha-smooth muscle actin; DAPI, 4’,6-diamidino-2-phenylindole; EA-naps, ellagic acid-loaded human serum albumin nanoparticles; EVG, elastic Verhoeff-Van Gieson; HE, hematoxylin-eosin; HFD, high-fat diet; LFD, low-fat diet; SEM, standard error of the mean.

### EA-naps increased LDLR expression and activated the EGFR-ERK signaling pathway in ApoE^-/-^ mice

We sought to test whether EA-naps can increase the expression of LDLR receptor and activate the EGFR-ERK signaling pathway *in vivo* as in the cell experiments. Therefore, we investigated the effects of EA-naps on LDLR expression and the EGFR-ERK signaling pathway in ApoE^-/-^ mice using western blotting and real-time quantitative polymerase chain reaction (PCR). The results showed that EA-naps significantly increased the LDLR mRNA and protein levels in the ApoE^-/-^ mouse liver (Figure 7A–B), and significantly increased the EGFR and ERK phosphorylation levels (Figure 7B). The in vivo effects were consistent with those observed in vitro.

**Figure 7.**
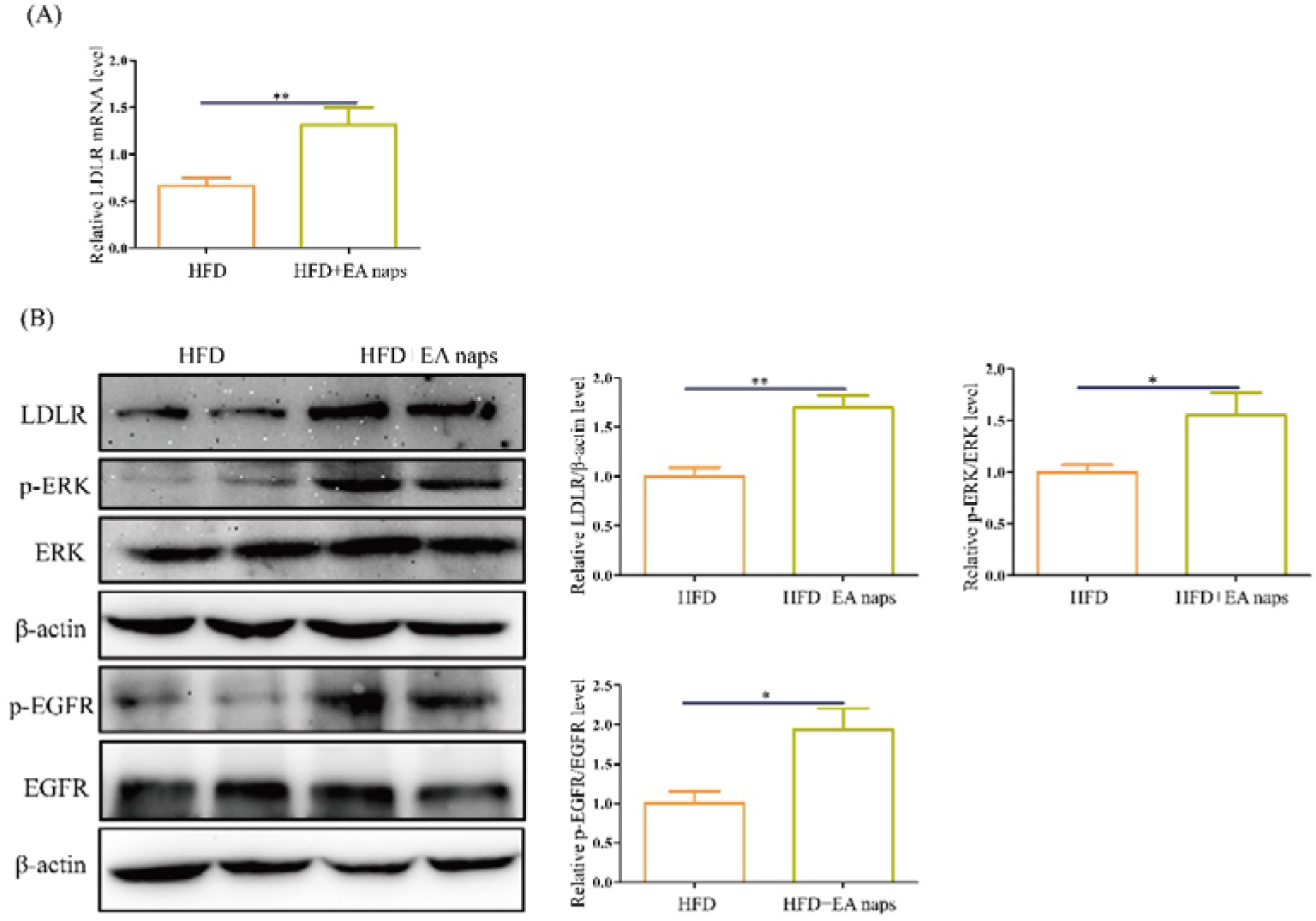
EA-naps activated the EGFR pathway to increase LDLR expression in HFD-fed ApoE^-/-^ mice. (A) mRNA levels of LDLR. (B) Protein levels of LDLR, p-ERK, and p-EGFR in the liver of ApoE^-/-^ mice. *P < 0.05, **P < 0.01, and ***P < 0.001. Data represent the mean ± SEM of six biological replicates. HFD group: high-fat and high-cholesterol diet, and treatment with uncoated ellagic acid nanoparticles; HFD+EA-naps group: high-fat and high-cholesterol diet, and treatment with coated ellagic acid nanoparticles. ApoE, apolipoprotein E; EA-naps, ellagic acid-loaded human serum albumin nanoparticles; EGFR, epidermal growth factor receptor; ERK, extracellular signal-regulated kinase; HFD, high-fat diet; IHC, immunohistochemistry; LDLR, low-density lipoprotein receptor; p-EGFR, phosphorylated-EGFR; p-ERK, phosphorylated-ERK; SEM, standard error of the mean.

### EA-naps improved blood lipid biochemical indices and reduced lipid accumulation in the liver

Abnormal lipid levels in blood is an important risk factor for the development of atherosclerosis. According to the blood lipid index, EA-naps significantly reduced the levels of TC (P < 0.05, Figure. 8A) and LDL-c (P < 0.05, Figure. 8C) in HFD-fed ApoE^-/-^ mice. However, its effect on triglycerides was not significant (P > 0.05, Figure 8B). Moreover, the comparison of liver appearance and oil Red O staining revealed that EA-naps also alleviated the HFD-induced lipid accumulation in the liver of ApoE^-/-^ mice (Figure 8D, E).

**Figure 8.**
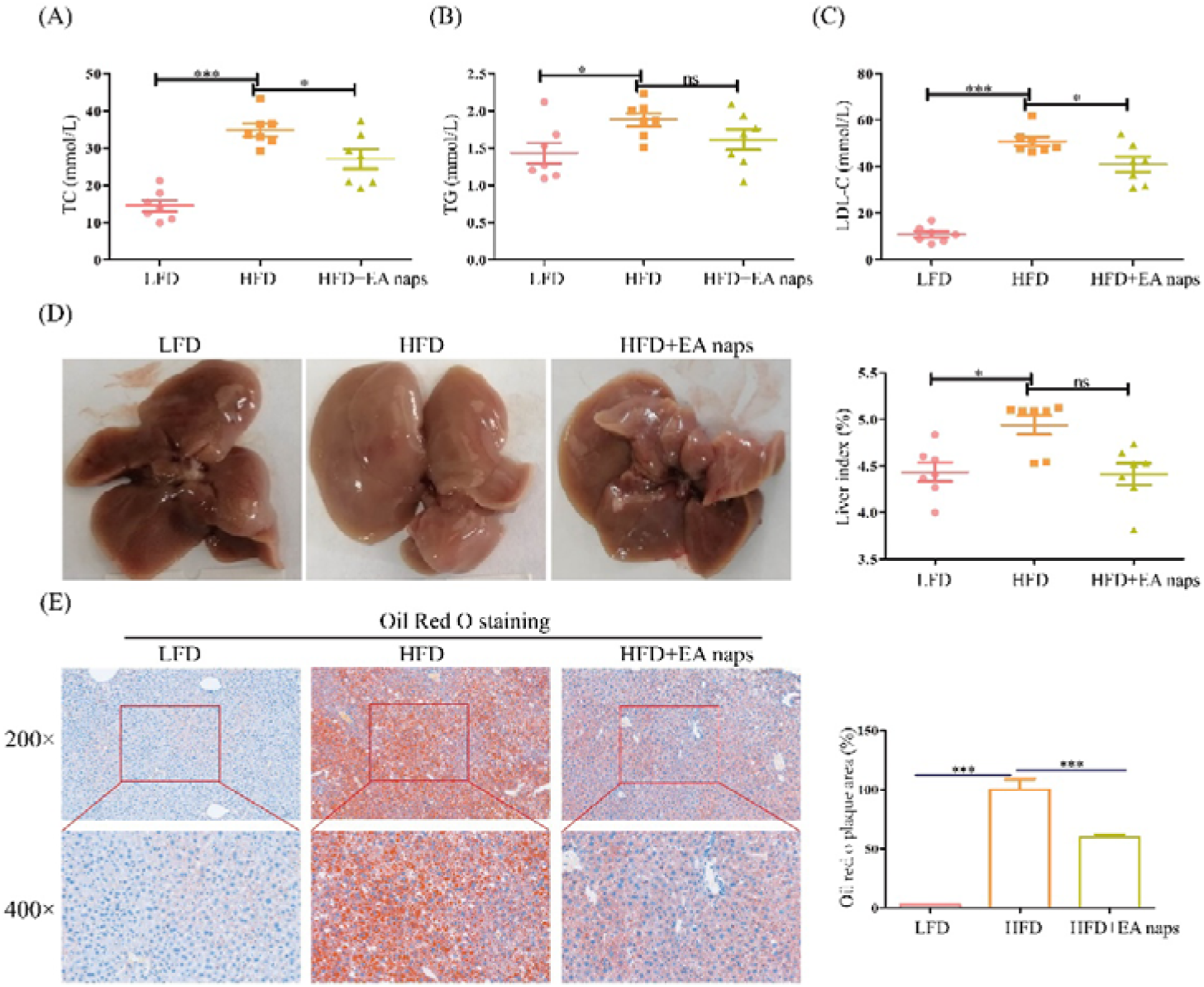
EA-naps improved blood biochemical parameters and liver lipid accumulation in HFD-fed ApoE^-/-^ mice. Levels of (A) TC, (B) TG, and (C) LDL-c in the serum of ApoE^-/-^ mice. (D) Livers of ApoE^-/-^ mice liver, and analysis of the liver index. (E) Livers of ApoE^-/-^ mice stained with oil Red O. *P < 0.05, **P < 0.01, and ***P < 0.001. Data represent the mean ± SEM of seven biological replicates. LFD group: chow diet; HFD group: high-fat and high-cholesterol diet, and treatment with uncoated ellagic acid nanoparticles; HFD+EA-naps: high-fat and high-cholesterol diet, and treatment with coated ellagic acid nanoparticles. ApoE, apolipoprotein E; EA-naps, ellagic acid-loaded human serum albumin nanoparticles; HFD, high-fat diet; LDL-c, low-density lipoprotein cholesterol; LFD, low-fat diet; SEM, standard error of the mean; TC, total cholesterol; TG, triglycerides.

## DISCUSSION

Atherosclerotic cardiovascular disease remains an important cause of death worldwide.^29^ Abnormally high LDL levels are a key risk factor for the development of atherosclerosis. LDLR can lower LDL levels by internalizing it from the blood into cells and mediating its degradation.^30,31^ Therefore, the promotion of LDLR protein expression through induction has been recognized as a key approach to the treatment of atherosclerosis. Previous studies have shown that EA induces the upregulation of LDLR in HepG2 cells through activation of the ERK signaling pathway.^32^ Nevertheless, the mechanism through which EA activates the ERK signaling pathway has not been revealed thus far.

The present study demonstrated that EA could enhance the mRNA stability of LDLR by activating the EGFR-ERK signaling pathway, induce an increase in the expression of LDLR mRNA and protein, and promote the uptake of LDL-c by HepG2 cells. This is similar to the mechanism through which 5-Azac increases the stability of LDLR mRNA by activating the ERK signaling pathway.^12^ However, unlike EA, 5-Azac causes the sustained activation of the inositol demand enzyme 1α (IRE1α) kinase domain and c-Jun N-terminal kinase (JNK), leading to the activation of the ERK signaling pathway and stabilization of LDLR mRNA. Our results showed that EA can directly bind to the extracellular segment of EGFR through hydrogen bonding and hydrophobic interaction, leading to activation of the EGFR-ERK signaling pathway, thus enhancing the stability of LDLR mRNA.

The potential activity of EA has been widely reported *in vitro*. However, its efficient delivery *in vivo* remains a challenge.^14^ Due to its low bioavailability and degradation in the digestive tract, EA cannot be transported in the body in a complete form and exert its effects.^33^ Therefore, it is difficult to determine the *in vivo* effects of EA on alleviating atherosclerosis; in addition, the mechanism of action of EA remains unclear. Numerous studies were conducted to resolve the problems of low solubility and bioavailability of EA. For example, Zhang et al.^34^ used protic solvent ethanol as the reaction solvent to prepare an EA-phospholipid complex; however, the recombination rate was low. Baghel et al.^35^ evaluated the ability of cellulose derivatives (e.g., carboxymethyl cellulose butyrate acetate, cellulose adipate propionate, and hydroxypropyl methyl cellulose acetate succinate) to form amorphous solid dispersions with EA by solvent evaporation for improving its solubility; nevertheless, the produced polymers were unstable. Savic et al.^36^ prepared EA inclusion complexes with β-cyclodextrin and hydroxypropyl β-cyclodextrin. However, the complexation of EA with cyclodextrin has been less successful in improving solubility. Moreover, cyclodextrin-based drugs are relatively expensive.

To overcome the challenge of *in vivo* EA delivery, we used HSA as a delivery carrier of hydrophobic drugs and prepared EA-naps through the self-assembly method.^37^ In the preparation method, the hydrophobicity of HSA molecules is increased by the addition of dithiothreitol (DTT). Subsequently, EA binds to the hydrophobic domain of HSA to mediate its encapsulation. The particle size of EA-naps prepared by in this study was 218.5 nm, which was suitable for intravenous administration. Of note, the formula exhibited a high encapsulation rate and stability (Zeta potential: −26.4 mV).

We also investigated whether EA-naps retain the properties of EA. Using cell experiments, we demonstrated that EA-naps can activate the EGFR signaling pathway and increase the levels of LDLR in HepG2 cells. EA-naps significantly improved atherosclerotic indicators in ApoE^-/-^ mice. Consistent with the *in vitro* analyses, *in vivo*, EA-naps also activated the EGFR-ERK signaling pathway and enhanced LDLR mRNA and protein expression.

Control of mRNA stability is a key regulatory mechanism for gene expression, and mRNA stability is controlled by a 3’ untranslated region containing AREs.^38^ It has been shown that some ARE-binding proteins (AUBPs) regulate the levels of ARE-containing mRNAs. For example, the RNA-binding protein human antigen R (HuR), which is commonly expressed in the *Drosophila* embryo fatal abnormal vision family, binds to AREs and stabilizes Mrna.^39^ In contrast, a number of other AUBPs, including tristetraprolin (TTP), butyrate response factor 1 (BRF1), and KH-type splicing regulatory protein (KHSRP), destabilize their target ARE-mRNAs.^40,41^ The results of our study demonstrated that EA can enhance the stability of LDLR mRNA at the post-transcriptional level by activating the EGFR-ERK signaling pathway. Nonetheless, further studies are warranted to investigate the precise mechanism underlying this post-transcriptional regulation, and identify the functional proteins involved in the process of enhancing the stability of LDLR mRNA.

We explored the binding pattern of EA and EGFR by molecular docking analysis. In general, the binding of ligands to polar and non-polar amino acid residues in the active site of the receptor generates hydrogen bonds and hydrophobic effects, respectively. Hydrogen bonds and hydrophobic effects are important modes of interaction between drug molecules and receptor proteins.^42,43^ Our results showed that EA forms hydrogen bonds and hydrophobic interactions with multiple amino acid residues of EGFR. These interactions may induce conformational changes in the extracellular domain of EGFR and ultimately lead to activation of the EGFR-ERK signaling pathway. Nevertheless, the specific molecular mechanisms, such as the specific binding sites of EA to EGFR and the series of effects associated with activation of the EGFR-ERK signaling pathway, remain to be further investigated. In addition, EGFR is often regarded as one of the proto-oncogenes involved in cell growth, cell signal transduction, and angiogenesis. Whether the effect of EA on the EGFR-ERK signaling pathway leads to imbalances in other cellular processes warrants further investigation.

## CONCLUSION

The present findings indicated that EA directly binds to the extracellular domain of EGFR, thus activating the EGFR-ERK signaling pathway, stabilizing LDLR mRNA, and promoting LDLR protein expression. Moreover, EA-naps prepared through the self-assembly method enabled intravenous administration in animals. The results of these *in vivo* experiments verified the effects of EA, namely the activation of the EGFR-ERK signaling pathway, increase in the levels of LDLR, and improvement of atherosclerosis-related indices. The results of this study may provide insight into the role of EA in the prevention and treatment of atherosclerosis. In addition, they may suggest new directions for the discovery of natural drugs against atherosclerosis.

## Acknowledgments

This study was supported by the Key Project of the International Cooperation Research Center for Eco-Friendly Food of Yunnan Province (2019ZG00904 and 2019ZG00909) and the Young talents support plan of Yunnan Province (YNWR-QNBJ-2018-083 and 2019HB021).

## Author contributions

Xuan-Jun Wang and Jun Sheng participated in the acquisition of resources and funds. Xuan-Jun Wang, Jun Sheng, Ye-Wei Huang participated in the conception, review and editing. Ye-Wei Huang and Li-Tian Wang are responsible for research, formal analysis, visualization, project management and writing of original manuscripts. Huai-Liu Yin and Dan-Dan Hu conducted investigations, data collation and verification.

